# Muscle Preflex Response to Perturbations in locomotion: In-vitro experiments and simulations with realistic boundary conditions

**DOI:** 10.1101/2023.02.15.528662

**Authors:** Matthew Araz, Sven Weidner, Fabio Izzi, Alexander Badri-Spröwitz, Tobias Siebert, Daniel F. B. Haeufle

## Abstract

Neuromuscular control loops feature substantial communication delays, but mammals run robustly even in the most adverse conditions. In-vivo experiments and computer simulation results suggest that muscles’ preflex—an immediate mechanical response to a perturbation—could be the critical contributor. Muscle preflexes act within a few milliseconds, an order of magnitude faster than neural reflexes. Their short-lasting activity makes mechanical preflexes hard to quantify in-vivo. Muscle models, on the other hand, require further improvement of their prediction accuracy during the non-standard conditions of perturbed locomotion. Additionally, muscles mechanically adapt by increased damping force. Our study aims to quantify the mechanical preflex work and test its mechanical force adaptation. We performed in-vitro experiments with biological muscle fibers under physiological boundary conditions, which we determined in computer simulations of perturbed hopping. Our findings show that muscles initially resist impacts with a stereotypical stiffness response—identified as short-range stiffness—regardless of the exact perturbation condition. We then observe a velocity adaptation to the force related to the amount of perturbation. The main contributor to the preflex work adaptation is not the force difference but the muscle fiber stretch difference. We find that both muscle stiffness and damping are activity-dependent properties. These results indicate that neural control could tune the preflex properties of muscles in expectation of ground conditions leading to previously inexplicable neuromuscular adaptation speeds.

## 2 Introduction

Legged locomotion in uneven terrain is a complex motorcontrol task performed seemingly effortless by humans and other animals (Blickhan et al. 2007). Animals reject unexpected ground perturbations(Daley and Biewener 2006; Müller et al. 2014), despite considerable sensorimotor transmission delays affecting the feedback control (More and Donelan 2018; More et al. 2010). This ability has long puzzled researchers in biomechanics and motorcontrol science. In-vivo research on perturbed legged locomotion suggests that the intrinsic mechanical properties of muscles are essential for motion stability during the first 30ms to 50ms after touchdown (Daley et al. 2009; Gordon et al. 2020; Nishikawa et al. 2007). During this brief interval, muscles and tendons react instantly through elastic and viscous-like properties. Brown and Loeb (2000, p. 161) labeled it *preflex*; the “…zero-delay, intrinsic response of a neuromusculoskeletal system to a perturbation”.

In-vivo walking experiments are essential for understanding robust locomotion. However, the functional mechanical and control coupling of muscle groups during whole-body movement complicates unveiling the regulatory principles behind preflexes, reflexes, and voluntary neuromuscular control. By artificially contracting individual muscle fibers, in-vitro research allows for precise isolation and investigation of muscles’ mechanical properties (Weidner et al. 2022). So far, a wide range of contraction settings have been explored, such as isometric, isotonic, and isovelocity(Brown et al. 2003; Gilliver et al. 2011; Tomalka et al. 2020). Yet, the exact boundary conditions of physiological muscle contraction are hard to replicate in in-vitro experiments. Cyclic fiber contractions during in-vitro experiments and follow-up work loop analyses are relatively realistic (Josephson 1985). So far, it is technically challenging to extract physiological trajectories of muscle contraction from in-vivo trials. Therefore, stretch-shortening cycle investigations that are limited to sinusoidal length trajectories (Darby et al. 2013) only roughly present locomotion conditions.

Previous simulation studies support the hypothesis that intrinsic muscle properties play a crucial role in stabilizing locomotion against disturbances (Gerritsen et al. 1998; John et al. 2013; Van Der Krogt et al. 2009). Simulation studies revealed that feedforward adjustment of muscle stimulation, as observed during human locomotion (Müller et al. 2015), may allow to adjust muscle mechanics according to perturbed impact conditions (Haeufle et al. 2018; Haeufle et al. 2010). As a means of investigation, computer simulations combine the advantages of in-vivo and in-vitro investigations. They enable the analysis of complex whole-body movements similar to in-vivo research while providing access to difficult-to-measure variables, similar to in-vitro experiments. Nevertheless, computer models depend on simplified assumptions. Most investigations of muscle preflex use classic Hill-type muscle models, which are restricted in their ability to describe muscle contraction outside of specific conditions (Siebert et al. 2021). Hill-type muscle models are parameterized with empirical data from isometric, isotonic, and isovelocity muscle fiber experiments, mostly at maximum activity, which are controlled experimental conditions differing greatly from in-vivo muscle loading. Compared to data from gait recordings, Hill-type muscle models were inaccurate in predicting muscle force during high-speed locomotion (Dick et al. 2017; Lee et al. 2013). Therefore, it still needs to be discovered to what extent simulation studies with Hill-type models can validate experimental research on muscle preflex. On the other hand, in-vitro experiments are required to test individual muscle fibers’ response to unexpected perturbation.

This study aims at understanding how individual muscle fibers exploit their intrinsic mechanical properties to respond to perturbations in realistic settings. We focus on how muscles’ elastic and viscous properties regulate energy absorption during the preflex phase to reject perturbations during locomotion impacts. We hypothesize that (1) muscles adapt their response according to the stretch velocities induced by ground perturbations, and (2) mechanical muscle properties can be tuned by changing activity level in advance to touch-down. We conducted muscle fiber experiments with realistic kinematic trajectories to prove our hypotheses. We obtained them by simulating vertical hopping driven by a Hill-type muscle model, at three levels of muscle activity and under three levels—step-up, no step, and step-down perturbations. Further, we derived a quasistatic-scenario for muscle fiber experiments with the same lengthening patterns over a much larger time to eliminate fiber’s velocity effect on muscle force production. These quasistatic-scenario experiments permit the separation of the elastic response from the viscous response of the muscle fibers. Finally, we compared simulations with muscle fiber experiments to test the accuracy of Hill-type models in explaining fiber response. Our results show that during the preflex phase, intrinsic muscle characteristics adjust the muscle force in response to the perturbation level. Our findings corroborate that muscle activity can tune mechanical muscle properties in advance. On the other hand, we found that Hill-type muscle models require modifications to mimic the muscle force response accurately.

## 3 Methods

The goal of this study was to test the force response of muscle fibers in realistic perturbation scenarios. Boundary conditions for in-vitro muscle fiber experiments were derived from a hopping simulation (Fig. 1A). We performed three simulations with a single-leg hopping model: a no-perturbation reference hopping (P0), a step-up perturbation (P↑), and a step-down perturbation (P↓). Fig. 1B shows the activity profile applied during the hopping simulation. During the first 30ms (preflex), the stimulation is kept constant due to the delay of neural transmission, and response against the perturbation is dependent only on elastic and viscous intrinsic properties. Then, stimulation rises linearly with time. We extracted kinematic trajectories and the activity state during the preflex phase (Fig. 1B) of the contractile element. The resulting data were used as boundary conditions for muscle fiber experiments and their corresponding simulations of the isolated contractile element (Fig. 1C-D). The highlighted preflex phase (Fig. 1B, shaded area)—is the focus of our study. Fig. 1B shows also the behavior shortly after the preflex phase. However, the data after the preflex is measured with constant activity levels, as the in-vitro setup did not allow for a time-controlled activity change. In the hopping simulation, the muscle activity rose after the preflex phase. We record force-length traces during these experiments and matching simulations and analyze preflex work-loops from the mechanical work of the muscle fiber. The following sections provide details of the experiments and simulations conducted.

**Figure 1:**
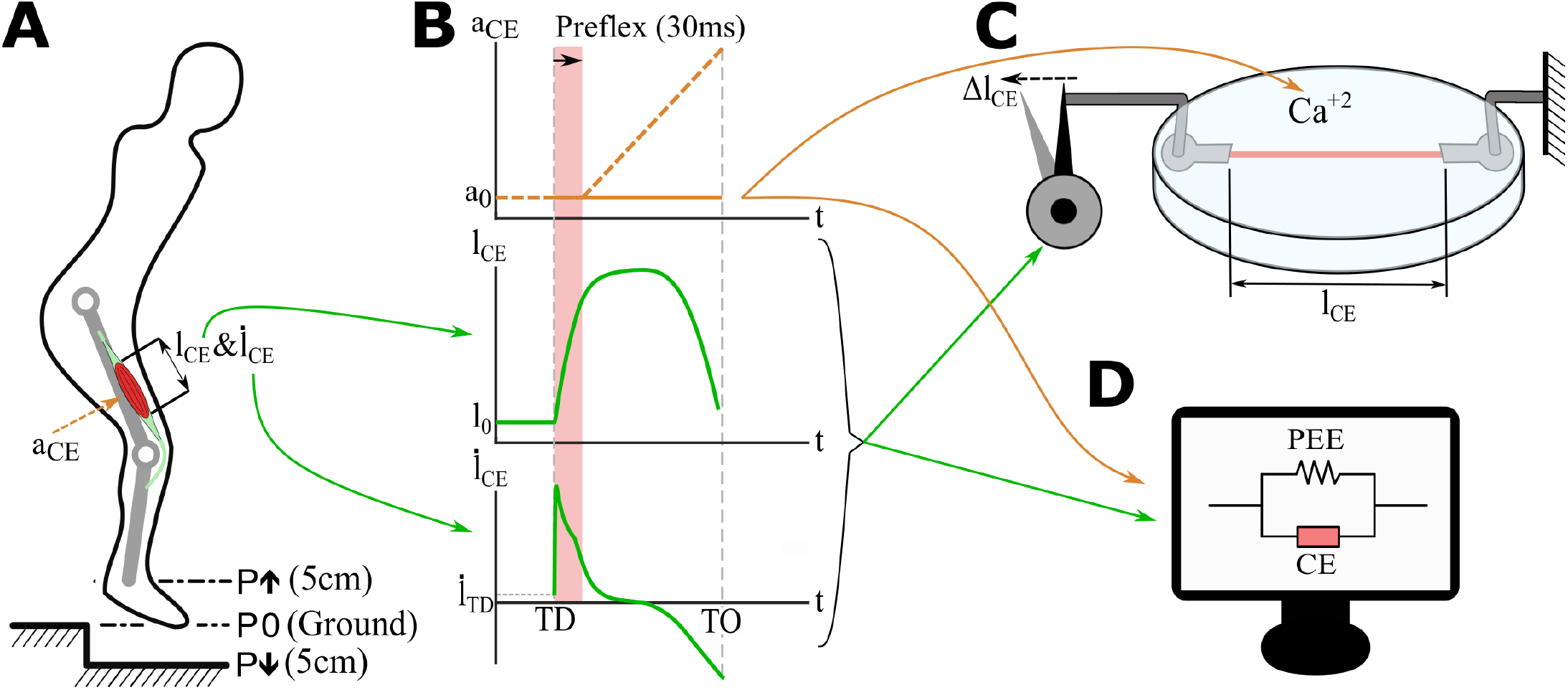
Simulations and in-vitro experiments applied in this study. A Hill-type model muscle-tendon unit drives a knee extensor unit of the hopper (Geyer et al. 2003; Haeufle et al. 2014; Izzi et al. 2022). Single-leg hopping is computer-simulated (**A**) with three perturbation scenarios: 5cm step-up (P↑), no perturbation (P0), and 5 cm step-down (P↓). The model outputs are length changes of the contractile element (l_CE_, **B**), and contraction velocities (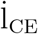, **B**). One hopping cycle from touch-down (TD) to toe-off (TO) is extracted for the analysis. Constant stimulation profile (a_CE_-orange line is shown only for 15% activity level, **B**) is then applied to in-vitro muscle fiber experiments (**C**) and isolated contractile element (CE) simulations (**D**). Constant stimulation in experiments and isolated contractile element simulations were used since the in-vitro setup does not allow changing stimulation levels within a stretch-shortening cycle. Since the muscle fibres used in in-vitro experiments (**C**) are isolated from the tendon, the contractile element (CE) and parallel elastic element (PEE) of the Hill-type muscle model are also isolated from the tendon unit (**D**).

### 3.1 Muscle Fiber Experiments

#### 3.1.1 Fiber Preparation

One M. extensor digitorum longus (*EDL*) was extracted from a single female Wistar rat, which was sacrificed with an overdose of CO_2_ shortly before. The specimen’s age was 8 to 10 months, at a body weight of 300 g to 350 g. The specimen was kept at a 12 h light and 12 h dark cycle at a housing temperature of 22 °C. The EDL muscle was obtained from the left hind limb. The experiment was conducted according to the guidelines of the Declaration of Helsinki and approved according to the German animal protection law (Tierschutzgesetz §4(3), permit no. T170/18ST).

The techniques used for muscle preparation, storage, and activation of skinned single muscle fibers were carried out as described in detail in Tomalka et al. (2017). Summarily, the EDL was prepared in 6 to 8 small fiber bundles, which were permeabilized in a skinning solution (see “Solutions”; Sec. 3.1.3) at 4 °C temperature directly after preparation. Fiber bundles were transferred to a storage solution made of 50% glycerol and 50% skinning solution, and kept at −20 °C for 6 to 8 weeks. Prior to conducting experiments, fibers were removed from the bundle using a dissecting microscope and fine forceps. Single fibers were cut to a length of 1mm. Aluminum T-shaped clips were folded around both ends of the fiber. The fiber was then treated with a skinning solution consisting of a relaxing solution with 1% vol/vol Triton-X 100 for 3min at 4 °C until the complete removal of internal fiber membranes (Linari et al. 2007).

#### 3.1.2 Experimental Setup

The skinned muscle fiber was transferred from the skinning solution to the experimental chamber of the fiber test apparatus (600A, Aurora Scientific, ON, Canada). One clipped end was attached to a length controller (model 308B, Aurora Scientific, ON, Canada) and the other end to a force transducer (Model 403A, Aurora Scientific, ON, Canada). Both attached ends were fixed with fingernail polish diluted with acetone (Getz et al. 1998). Transitions from the fiber end to the clip were treated with glutaraldehyde in rigor solution to improve mechanical performance and stability during the experiment (Hilber and Galler 1998).

The central fiber segment was focused and used to measure the sarcomere length (Weidner et al. 2022), which was set to 2.5 μm (means ± standard deviation) in the beginning. At this optimal sarcomere length l_s0_ the fiber produces its maximum force F_0_ (Stephenson and Williams 1982). The corresponding muscle fiber length is defined as l_opt_. The height (h) and width (w) of the fiber was measured in 0.1mm increments over the entire length of the fiber using a 10× extra long working distance dry lens (NA 0.60, Nikon, Japan) and a 10× eyepiece. The cross-sectional area of the muscle fiber was determined 5.25e–9m^2^ (±1.5e–9) assuming an elliptical cross-section (π × *h* × w/4).

A high-speed video system (Aurora Scientific, 901B, Canada) in combination with a 10x extra long working distance dry objective (NA 0.40, Nikon, Japan) and an accessory lens (2.5×, Nikon, Japan) visualized and tracked dynamic changes in the sarcomere length. Videos were recorded at 300 Hz recording frequency.

#### 3.1.3 Solutions

The relaxing solution contained 0.1mol TES, 7.7mmol MgCl_2_, 5.44mmol Na_2_ATP Na_2_ATP, 25mmol EGTA, 19.11 mmol Na_2_CP, 10mmol GLH (pCa 9.0). The preactivating solution contained 0.1mol TES, 6.93mmol MgCl_2_, 5.45mmol Na_2_ATP, 0.1mmol EGTA, 19.49mmol Na_2_CP, 10mmol GLH, and 24.9mmol HDTA. The skinning solution contained 0.17mol potassium propionate, 2.5mmol MgCl_2_, 2.5mmol Na_2_ATP, 5mmol EGTA, 10mmol IMID, and 0.2mmol PMSF. Recipes for activation solutions (‘ACT’) are shown in Tab. 1. The storage solution is the same as the skinning solution, except for the presence of 10 mmol GLH and 50% glycerol vol/vol. Cysteine and cysteine/serine protease inhibitors [trans-epoxysuccinyl-L-leucylamido-(4-guanidino) butane, E-64, 10mM; leupeptin, 20mg/mL] were added to all solutions to preserve lattice proteins and thus sarcomere homogeneity (Linari et al. 2007; Tomalka et al. 2017). KOH was applied to adjust to a pH 7.1 at 12 °C. Then, 450 U/mL of creatine kinase were added to all except skinning and storage solutions. Creatine kinase was obtained from Roche, Mannheim, Germany, and the remaining chemicals were obtained from Sigma, St Louis, MO. According to our calibration curve (Fig. S1) we chose concentrations of 6.73 pCa, 6.34 pCa and 6.3 pCa to best match the simulations boundary conditions.

**Table 1:** Recipe of activation solutions used, values are in [mmol].

|  | 5 %ACT | 15 %ACT | 25 %ACT | 100 %ACT |
| --- | --- | --- | --- | --- |
| TES | 100 | 100 | 100 | 100 |
| MgCl<sub>2</sub> | 7.183 | 6.995 | 6.98 | 6.76 |
| EGTA | 11.25 | 6.25 | 5.85 | 0 |
| CaEGTA | 13.75 | 18.75 | 19.15 | 25 |
| Na<sub>2</sub>ATP | 5.451 | 5.455 | 5.455 | 5.46 |
| KPi | 0 | 0 | 0 | 0 |
| Na<sub>2</sub>CP | 19.319 | 19.395 | 19.4 | 19.49 |
| GSH | 10 | 10 | 10 | 10 |

#### 3.1.4 Experimental Protocol

All experimental trials were conducted at a solution temperature of 12 °C (±0.1). At this temperature, the skinned muscle fibers proved stable during work-loop experiments (Tomalka et al. 2020, 2021). Fibers can tolerate activation and active stretch protocols over a long period (Ranatunga 1984, 1982). A three-step approach was used to activate the fibers by calcium diffusion. First, muscle fibers were immersed for 60 s in a preactivation solution for equilibration. The fiber was then transferred to the activation solution. This led to a rapid increase in force until a plateau was reached. We defined the plateau as an isometric force increase of less than 1% rise of force within 1.5 s. After reaching the plateau, the perturbation was carried out. In the last step, the fiber was transferred to the relaxing solution, in which it was prepared for the subsequent activation for 400 s using cycling protocols (Tomalka et al. 2017).

The in-vitro experiment included isometric contractions at optimal fiber length and three hopping stretch-shortening cycles based on the simulation data of the hopping model (Sec. 3.2). First, the activity level of the fiber in three sub-maximal conditions was checked using isometric contractions in 5 %, 15 %, 25 %, and supra maximal activation solution at optimal fiber length. This step ensured matching boundary conditions with the simulation data. A flow chart of an experimental day for a single fiber is shown in Fig. S2. Hopping stretch-shortening cycles were applied to the fiber according to length and velocity data extracted from the simulation in the simulated dynamic-scenario, and a quasistatic-scenario (see Sec. 3.2 for more details). Both the order of stretch-shortening profiles and the order of activity levels within a perturbation, called a “block”, were randomized. Each block was surrounded by isometric activity at optimal fiber length and full activity to take into account fiber degradation during force data normalization (Fig. S2).

Velocity, force, and length data were recorded at 1 kHz for isometric and quasistatic-scenario trials and 10 kHz for high-speed trials with an A/D interface (604A, Aurora Scientific, ON, Canada). The data acquisition was carried out with real-time software (600A, Aurora Scientific, ON, Canada). Data were loaded into MATLAB (MathWorks, MA, USA) and analyzed with a custom-written script. Forces during perturbation trials for every single fiber were divided by individual F_0_, and likewise, fiber length l by individual l_opt_, to normalize them.

### 3.2 Simulations

#### 3.2.1 Generating Hopping Trajectories in Simulation

To identify realistic boundary conditions for in-vitro muscle fiber experiments, we extracted contractile element kinematics from a single-leg hopper simulation ((Izzi et al. 2022) based on the model by Geyer et al. (2003)). The single-leg hopper is driven by a Hill-type muscle-tendon unit (MTU) model. The model considers four elements: a contractile element representing the muscle fibers, a parallel elastic (PEE), a serial elastic (SEE), and a series damping (SDE) element (Haeufle et al. 2014). The modeled muscle-tendon unit extends the knee joint (Fig. 1A). The leg features two massless segments connected by the knee hinge joint. The body mass is represented as a point mass located at the hip joint.

We simulated stable periodic hopping with the hopper model and introduced a step-up and a stepdown perturbation (Fig. 1A). During the flight phase, the muscle is stimulated with 15% constant stimulation, and the knee joint is fixed. After touch-down, the constant stimulation level continued for 30ms throughout the preflex phase and then increased with a ramp input (*b* = 10s^-1^, Fig. 1B). Despite the constant stimulation during the preflex phase, the contractile element can change its force due to the elastic and viscous intrinsic properties, which are related to force-length and force-velocity, respectively. Since the Hill-type muscle is extending the knee in the hopping simulation, muscle-tendon unit and contractile element are stretched at the initial phases of the stretch-shortening cycle. Thus, the model operates at the eccentric section of the force-velocity relation (Haeufle et al. 2014):

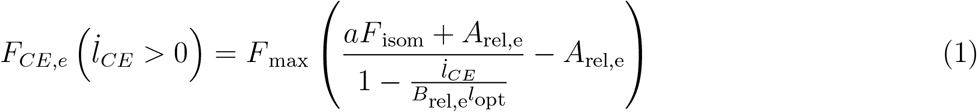

Here 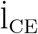 is the fiber contraction velocity, F_max_ is the maximum isometric force that the contractile element can generate, F_isom_ is the isometric force that the contractile element generates according to the current muscle length, A_rel,e_ and B_rel,e_ are the normalized Hill parameters for the eccentric phase, and *a* is the activity level. Even though the force-length relation represented in the Hill-type muscle model is independent of the current activity state, force-velocity relations are affected by activity. Fig. 2 shows force-velocity traces that the Hill-type muscle model generates at 5%, 15%, 25%, and full activity (100%) states.

**Figure 2:**
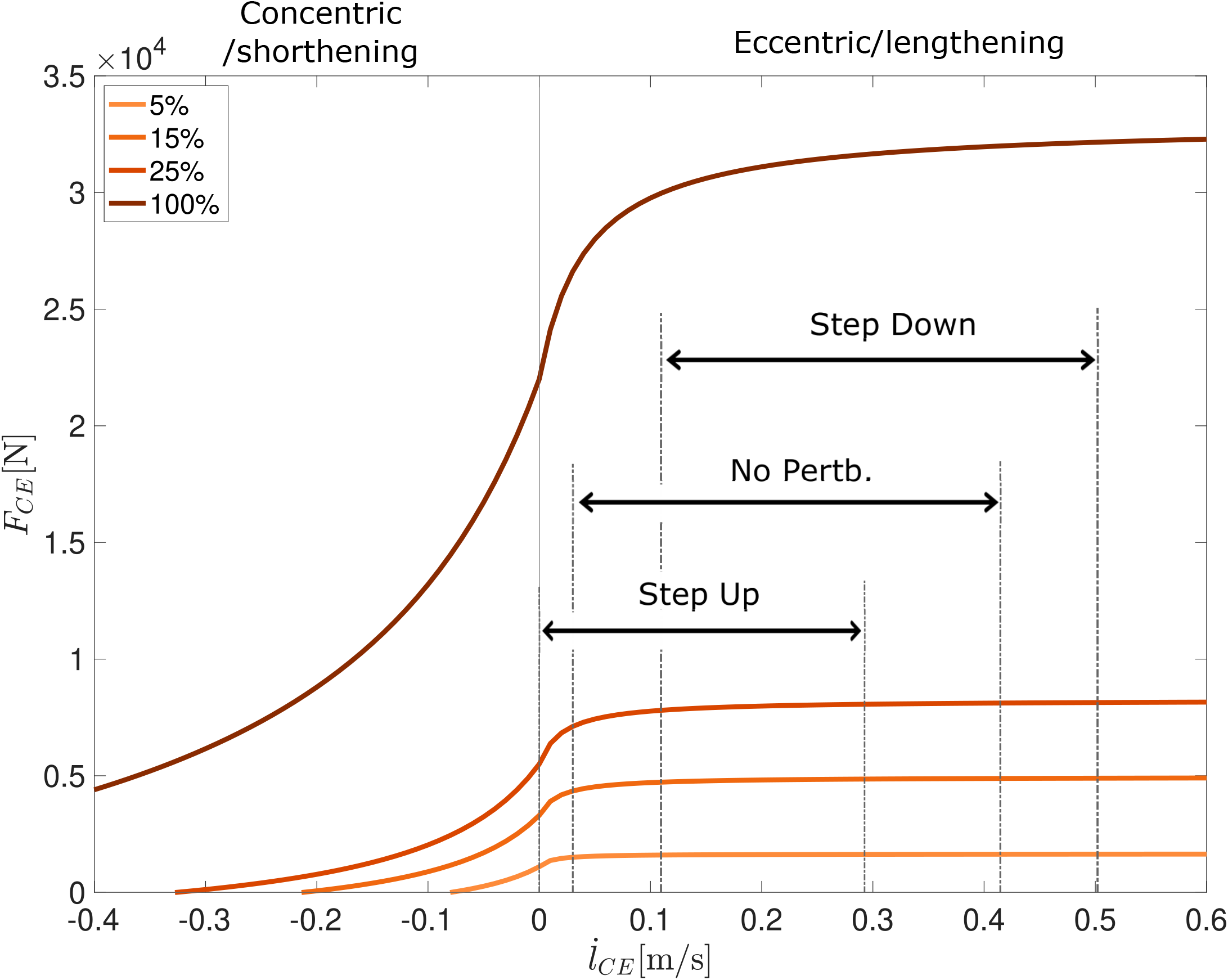
Force-Velocity relation for the contractile element for activity levels of 5%, 15%, 25% and 100%.

For a comparison between the biological muscle fiber and the Hill-type muscle model behavior, the parameters of the isometric force-length curve of the model were optimized to fit experimental data (Stephenson 2003). More precisely, the width of the normalized bell-curve ΔW_limb_ and its exponent *ν*_CE,limb_ of the ascending limb were optimized with the *lsqcurvefit* function (MATLAB 2021b).

#### 3.2.2 Extracting Boundary Conditions

We simulated hopping for no-perturbation locomotion on ground level, 5 cm step-up perturbation, and 5 cm step-down perturbation. Beyond step-down perturbations of 5 cm, the single-leg hopper generates unstable hopping patterns. Thus we decided to use a maximum perturbation height of 5 cm. The contractile element length and velocity profiles were extracted for each perturbation level. These kinematic data were used in muscle fiber experiments and isolated contractile element simulations to compare their reactions to the perturbations during the preflex phase.

We further derived quasistatic-scenario boundary conditions to differentiate between the muscle fibers’ velocity-dependent and length-dependent force adaptation. To create a length-dependent force adaptation, we generated quasistatic-scenario boundary conditions for each perturbation level. In these conditions, the time duration of the contractile element lengthening profiles obtained from each perturbation level was expanded by 80 times compared to the original duration. Thereby, the contraction velocity was decreased to negligible levels without exceeding the minimum speed limits of the experimental setup. Hence, the viscous contribution was eliminated from the muscle fiber force response, and the muscles only reacted with their elastic properties to the perturbations.

#### 3.2.3 Simulating Isolated Contractile Element Response

In the hopping simulation, the Hill-type muscle model calculates contractile element kinematics according to the dynamic balance of the serial (SEE & SDE) and contractile (CE & PEE) side of the model. However, in-vitro experiments are conducted only with isolated muscle fascicles. To match in-vitro conditions, we ran simulations solely with an isolated contractile element. Thereby, the isolated responses of the CE—corresponding to the muscle fiber—were calculated according to the provided contractile element length (l_CE_), contraction velocity 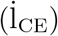 and activity (a):

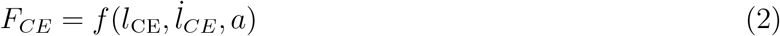

We obtained the kinematic data of contractile element from hopping simulations for step-up and stepdown perturbations. No perturbation cases were supplied as input to the isolated contractile element model with constant activity levels of 5%, 15% and 25%. Although the activity level increases after the preflex phase during hopping simulations, isolated contractile element model simulations must be kept constant to reproduce the conditions of in-vitro muscle fiber experiments. In the experimental setup, the stimulation level is arranged with chemical baths explained in Sec. 3.1.3. The setup allows only a single stimulation level for each stretch-shortening cycle. Therefore, stimulation levels were kept constant in isolated contractile element simulations to match the experimental in-vitro conditions.

### 3.3 Data Analysis and Statistics

#### 3.3.1 Data Analysis

The analysis of stretch-shortening cycles of both experimental and simulated fiber contractions focused on the preflex phase, which is the first 30ms of the dynamic-scenario scenarios. We analyzed data in the quasistatic-scenario scenario until the fiber lengthening reached the same level as at the end of the preflex phase in the dynamic-scenario scenario. We calculated the areas under force-length curves with the *trapz* function (MATLAB 2021b) to measure the work done by the muscle fiber and the isolated contractile element model. In addition, we estimated the muscle fiber’s short-range stiffness from the slope of fitted force-length curves during the initial phase of preflex (from 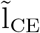 (l_CE_/l_opt_) 0.57 to 0.59). Then, to observe the effect of velocity on stiffness, we calculated the stiffness during the quasistatic-scenario stretch for the same boundary conditions.

#### 3.3.2 Statistics

SPSS 27 (IBM Corp., Armonk, NY) was used for the statistical analysis, with a significance level of *p* = 0.05. Initially, we tested for normal data distribution by running a Shapiro-Wilk, which showed negative. Hence, we used a Friedman test to elucidate differences between the applied perturbations within one activity level. We executed comparisons pairwise for post hoc experimental data analysis. Results were fed into a Bonferroni correction to take multiple testing into account. We tested for differences between similar activity levels and applied perturbation between dynamic-scenario and quasistatic-scenario conditions with a two-sample paired sign test. Effect sizes for the pairwise comparisons were classified as small (*d* < 0.3), medium (0.3 < *d* < 0.5), and large (*d* > 0.5) using Cohen’s *d* (Cohen 2013).

### 4 Results

During in-vitro experiments, we found that intrinsic muscle properties adapt the force response to the perturbation condition within the preflex phase (Fig. 3A, thick lines). Work loops of dynamic-scenario experiments for skinned fibers show muscle fibers are initially responding with similar force and with a linear and increasing trend between touch down and 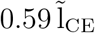 in all perturbations (Fig. 3A). After 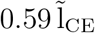, the force differs depending on the perturbation level, i.e., force is highest in the step-down perturbation (Fig. 3A).

**Figure 3:**
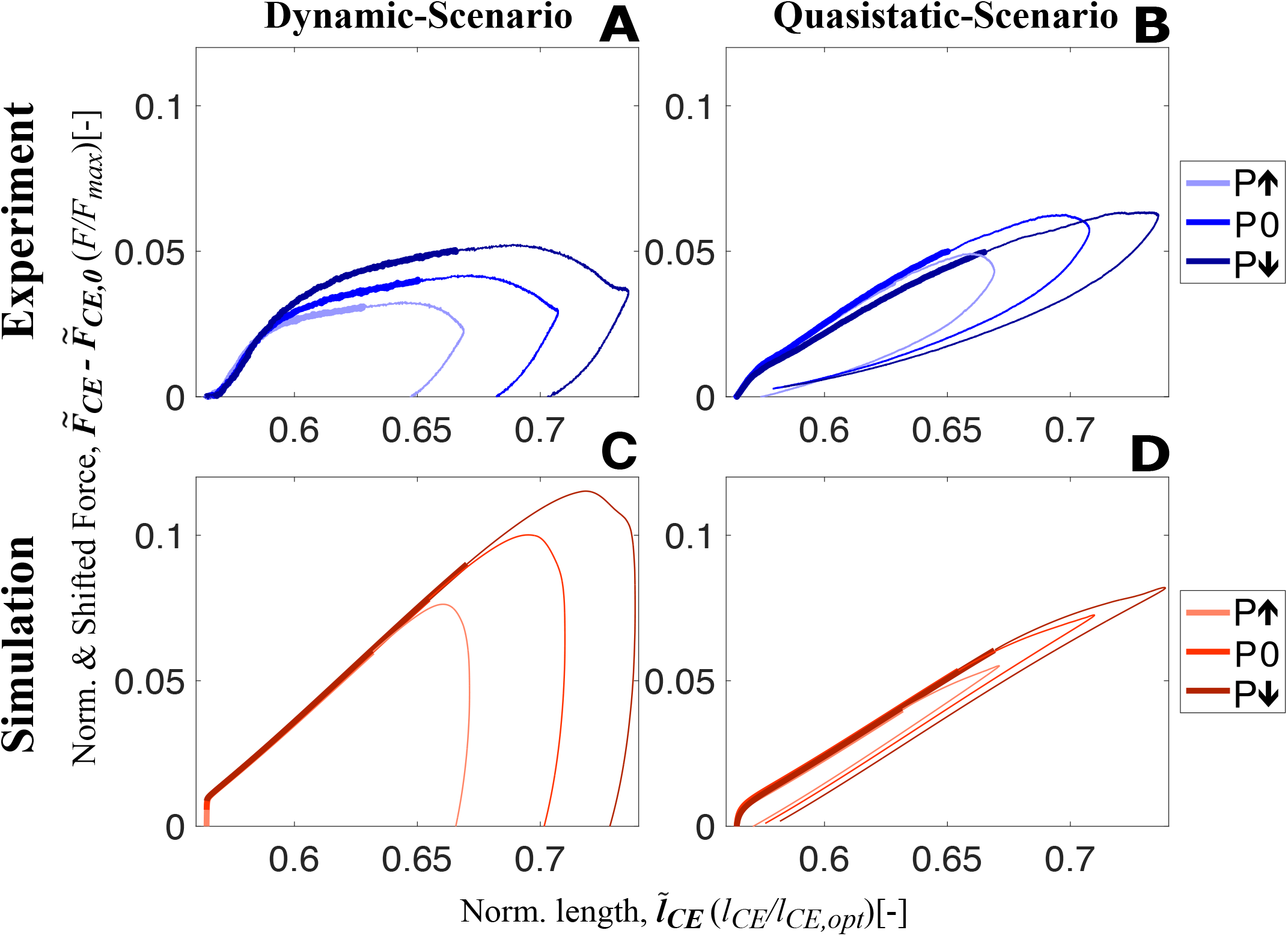
Shifted work loops for dynamic-scenario and quasistatic-scenario analysis step up(P↑), no (P0) and step down (P↓) perturbations for both experiments (**A-B**) and simulations (**C-D**) at 15% activity level (Work loops for 5% and 25% are shared in Figs. S3 and S4, respectively). The experimental data presented on **A** and **B** show the mean of all experimental trials. From touch-down to toe-off, all stretch-shortening cycle loops are plotted in the clockwise direction, and the thick and thin sections of the loops represent the preflex and remaining part of the stretch-shortening cycle, respectively. The preflex stretch gets longer from step-up to step-down perturbation since the muscle stretches faster in the same amount of time.

The response we observed from the skinned fiber experiments does not match with the prediction of the isolated contractile element element of the Hill-type muscle model (Fig. 3A,C). For the Hill-type muscle model, an effect of the perturbation is only observed at touchdown (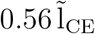, Fig. 3C). Right after touchdown, the response of the Hill-type muscle model reaches the same force level regardless of the perturbation state and then increases with the same linear trend. Therefore, the model did not predict the adaptation of the force magnitude to the perturbation condition.

Contrary to the dynamic-scenario experiments (Fig. 3A), the force response of skinned fibers in quasistatic-scenario stretches does not change according to the perturbation level (Fig. 3B), during preflex. Initial forces and the rise in force are similar for all perturbation levels. This result matches the prediction of the isolated contractile element element of the Hill-type muscle model (Fig. 3D).

We found that intrinsic muscle properties adapt the mechanical work during the preflex phase (preflex work) according to the perturbation condition. This is true in dynamic-scenario and quasistatic-scenario tests, experiments and simulation, and all activity levels (Fig. 4).

**Figure 4:**
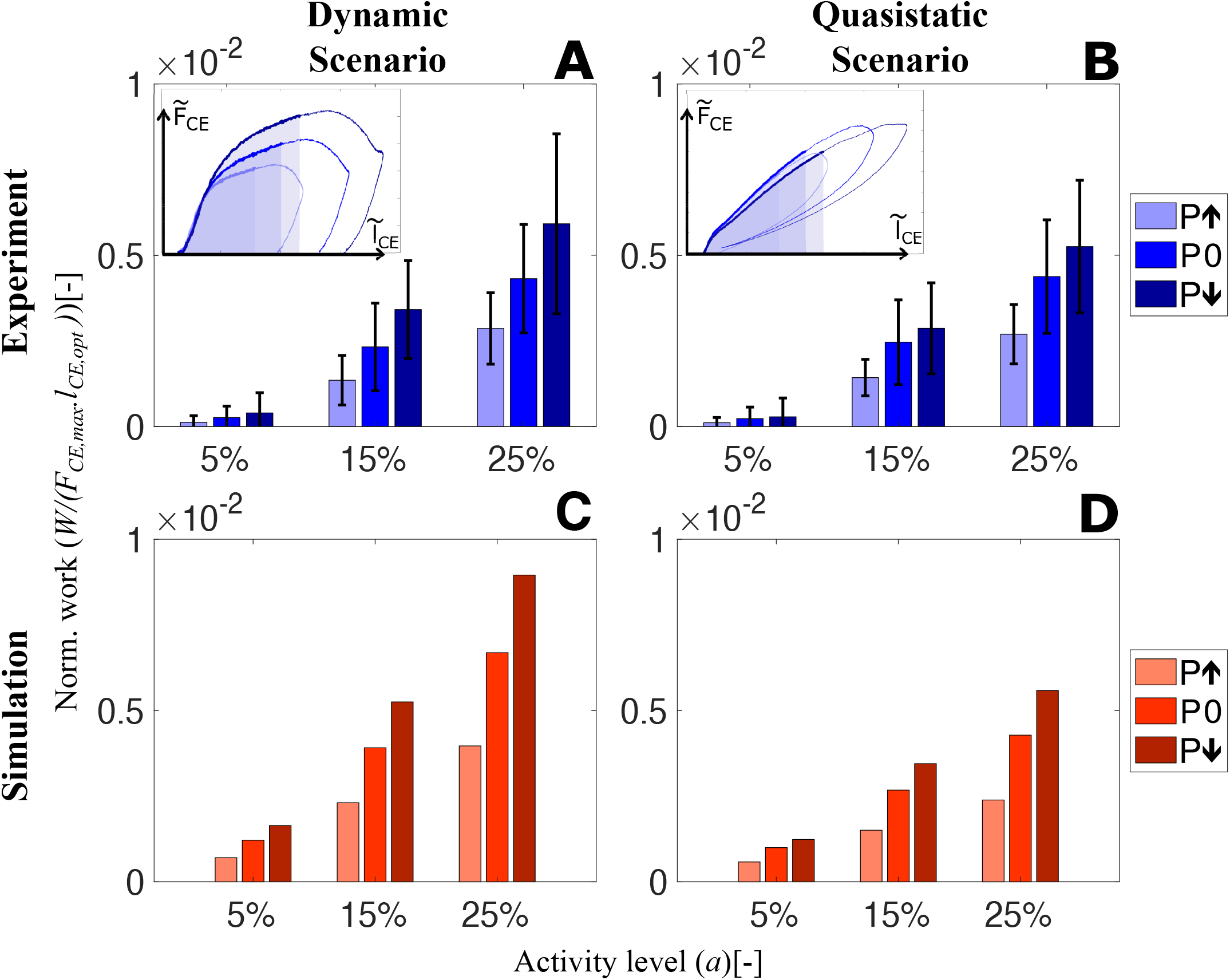
The amount of preflex work done by skinned fibers and the isolated contractile element of the Hill-type muscle model for all perturbation states and activity levels. In the dynamic-scenario analysis, work done during the preflex phase was calculated as the area under the force curve. Shaded areas in the insets (**A**) and (**B**) indicate changing perturbation levels. In the quasistatic-scenario analysis, the area till the lengthening reached at the end of the preflex phase was analyzed for each perturbation level. Bars in (**A-B**) and (**C-D**) indicate the work done by skinned muscle fibers and the isolated contractile element element at dynamic-scenario and quasistatic-scenario, respectively. We calculated and indicate the area of the normalized works loops (Fig. 3). Hence, there is no need to match the parameters of the hopping simulation to experimental muscle size.

The preflex work increases when comparing the step-up to the step-down perturbation, according to the dynamic-scenario analysis of skinned fibers (Fig. 4A). Albeit no significant difference among perturbation states at 5% activity level (*p* = 0.169), a perturbation influence on preflex work is observable for 15% (*χ*^2^ = 12.61; *p* = 0.002) and 25% (*χ*^2^ = 14; *p* = 0.001) activity levels (Tab. S1).

Preflex work changes significantly between activity levels. For the same kinematic profiles, the work done by skinned fibers increases if they are activated more (5% to 25% activity level; p = 0.001). The work differences between the perturbation conditions increase with an increase in activity level. See the supplementary materials (Tab. S2) for further details.

In the dynamic-scenario analysis, the muscle fibers’ response is a combination of two mechanical features; elasticity and viscosity. To identify their individual contributions, we minimized the parameter responsible for the viscous contribution—the stretch velocity. We performed quasistatic-scenario experiments, where muscles were stretched with the same lengthening profiles as in dynamic-scenario experiments, but at super-low velocities. Hence, with this experimental design, we expect to see only the elastic muscle fiber response. Still, even at negligible stretch velocities, we observed a similar preflex work trend between perturbation levels in quasistatic-scenario experiments and dynamic-scenario experiments (Fig. 4A,B).

Surprisingly, the Hill-type muscle model predicted the amount of preflex work accurately for the dynamic-scenario experiments (Fig. 4A,C) and the quasistatic-scenario experiments (Fig. 4B,D). Such correct prediction is an expected outcome for quasistatic-scenario experiments but not for dynamic-scenario experiments; the model’s dynamics differed compared to the muscle fiber dynamics (Fig. 3A,C).

Work loops obtained from the dynamic-scenario analysis (Fig. 3A, Fig. S3A, and Fig. S4A) show that the force response of the muscle fibers is almost identical within the short-range stiffness (Rack and Westbury 1974) regardless of the velocity profile. Only after the short-range stiffness phase, the force and energy are affected by velocity (Fig. 5). To understand the influence of velocity on the preflex work, we measure the area after the end of short-range stiffness phase until the end of preflex, for the step-up perturbation condition (Fig. 5A, inset: shaded areas). Work done in this phase is slightly higher for the faster stretches at 15% and 25% activity levels. However, the differences between the perturbation levels are not significant (15% activity: *χ*^2^ = 4.67, *p* = 0.097; 25% activity: *χ*^2^ = 3.56, *p* = 0.167; See Tab. S1 for more details).

**Figure 5:**
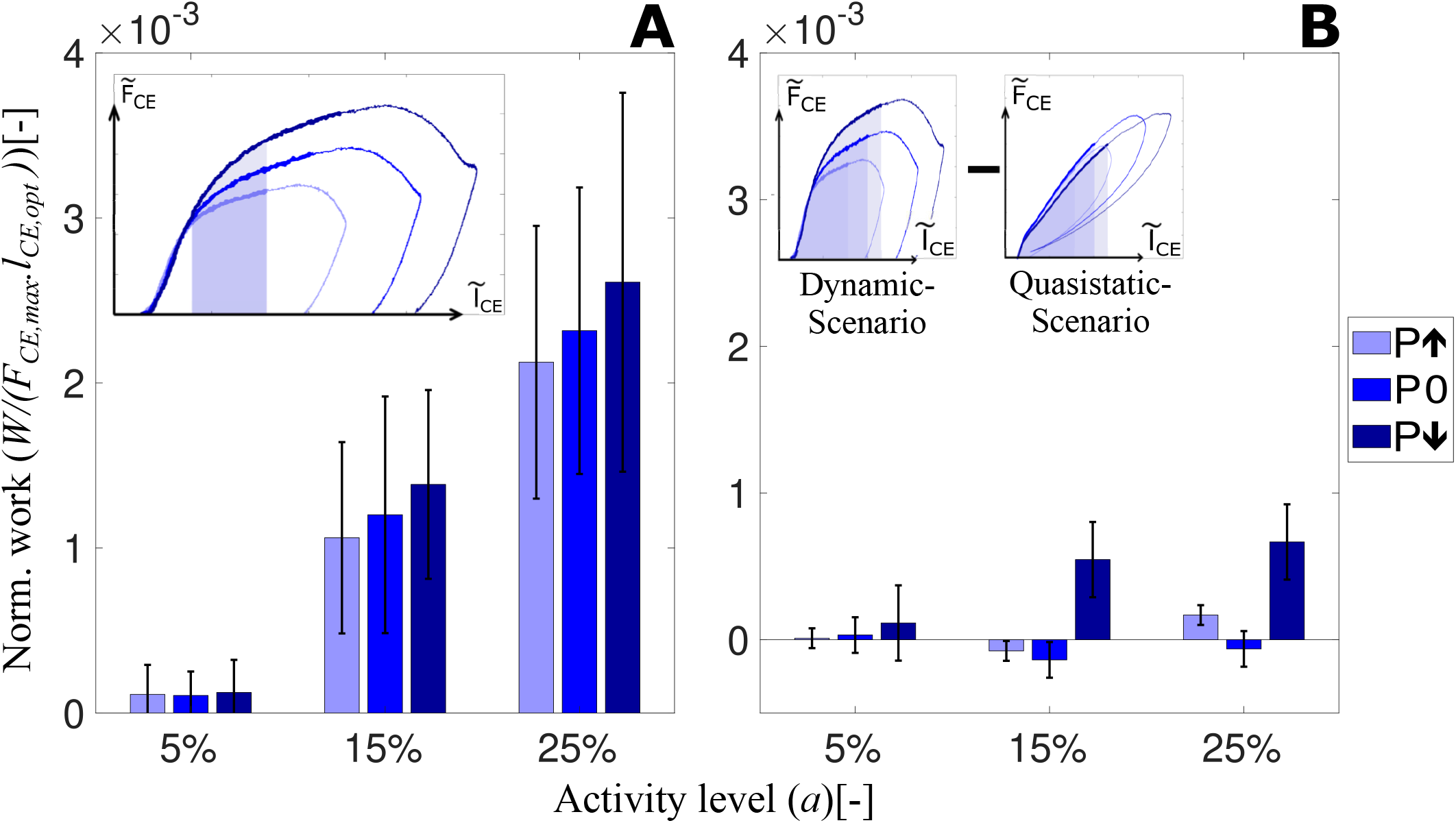
Influence of velocity adaptation on preflex work is represented. **A** shows the dissipated energies at dynamic-scenario experiments after the short-range stiffness till the end of the ‘step up’ perturbation phase (shaded area shown in the inset). Elastic contribution is kept equal for all perturbation states. Thus, the difference in energy will be caused by the difference in velocity profiles. **B** shows the preflex work difference between the dynamic-scenario and quasistatic-scenario experiments for each perturbation level. The preflex work is shown in the insets as a shaded area and for multiple conditions.

We compare activity levels effecting muscle work to see whether muscles’ viscous properties are tunable. Our results show that the activity level influences the amount of viscous contribution (Fig. 5A) similar to the preflex work (Fig. 4A). For the same kinematic profiles, a rising activity level causes a work increase by viscous characteristics of muscle fibers (15% to 25% activity level; *p* = 0.001).

To understand the velocity-related adaptation throughout the preflex phase, we subtract the work done in the quasistatic-scenario condition from dynamic-scenario experiments, shown as inset in Fig. 5B. Surprisingly, the amount of work done by muscle fibers at dynamic-scenario and quasistatic-scenario experiments are almost identical, and we measure no significant effect of the velocity on the preflex work (Tab. S3). Muscle fibers did more work in three conditions of the quasistatic-scenario case. So neither the velocity differences between perturbations nor between quasistatic-scenario and dynamic-scenario experiments had a relevant effect on the preflex work.

Analysis of the short-range stiffness shows no difference between perturbations but significant differences between activity levels (Fig. 6A). In quasistatic-scenario experiments, we find no significant differences between perturbation levels. However, short-range stiffness is smaller in quasistatic-scenario stretches (Fig. 6B) than in dynamic-scenario experiments (Fig. 6A), and the difference between them increases with higher activities (Tab. S3). Hence, short-range stiffness is increasing from quasistatic-scenario (Fig. 6D) to dynamic-scenario velocities (Fig. 6C). However, short-range stiffness does not change according to the difference in velocity between the perturbation levels (Fig. 6C, different shades of thick blue lines). Besides, the stiffness value can be arranged by changing the activity level.

**Figure 6:**
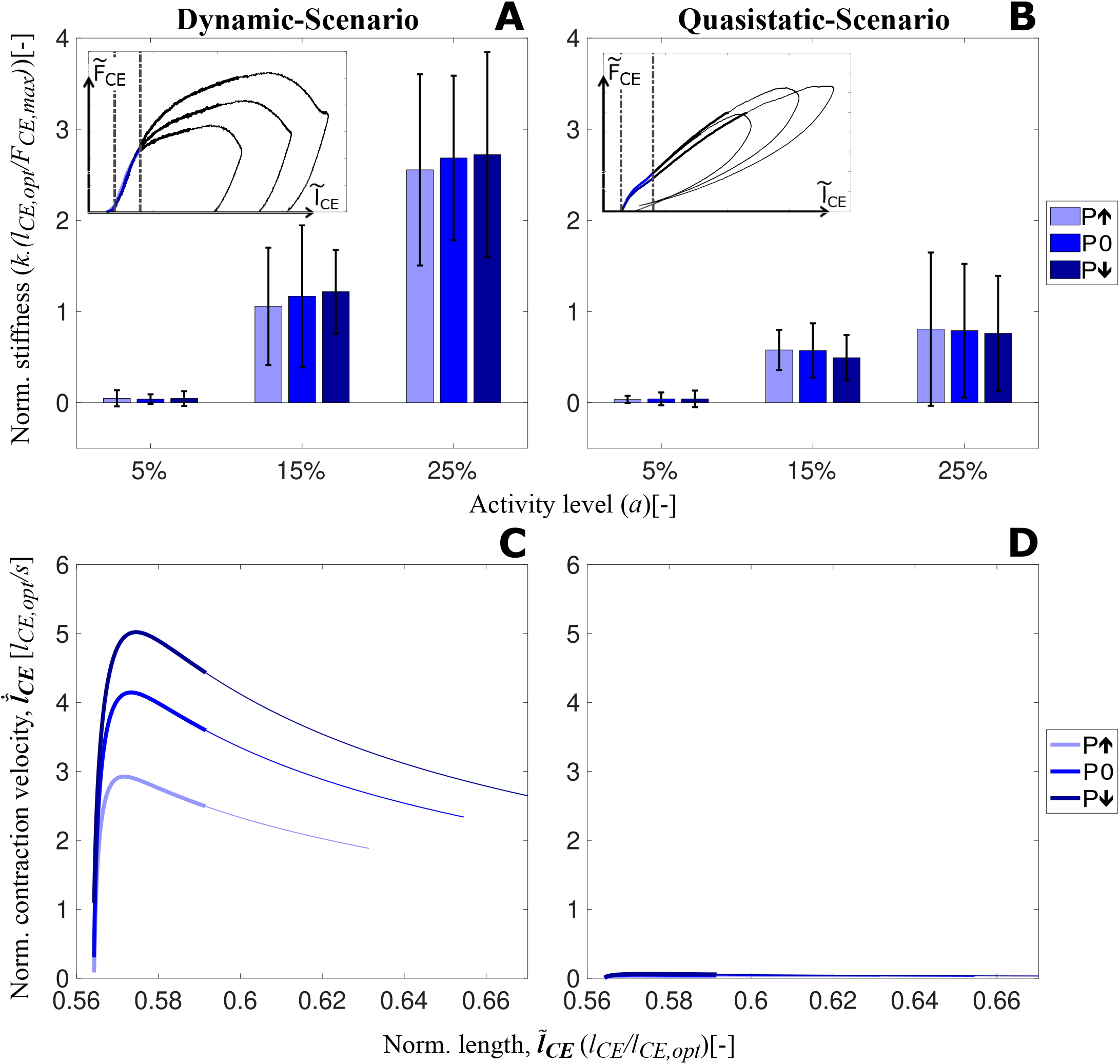
Short-range stiffness of muscle fibers during the dynamic-scenario (**A**). Stiffness amount during the short-range stiffness lengthening during quasistatic-scenario experiments (**B**) for all perturbation and activity levels are shown in the bar charts. Boundary conditions for the stiffness calculations for both speed conditions are shown as insets. In addition, velocity-length profiles during the preflex phase are presented in (**C**) and (**D**) for actual and quasistatic-scenario experiments, respectively. Thick lines show the short-range stiffness phase, and thin lines present the remaining part of the preflex phase.

## 5 Discussion

In this study, we presented the first in-vitro experiments conducted under realistic boundary conditions and activity levels of perturbed hopping. Our aim is to understand how an individual muscle fiber adapts its intrinsic properties to reject perturbation during locomotion. We extracted the boundary conditions from a hopping simulation for three levels of perturbations. Here, we discuss surprising outcomes that we observed from our in-vitro experiments and simulations. As hypothesized, muscles adapt their force response to the stretch velocities. However, this adaptation is not the main contributor to the preflex work. In addition, we observed that muscle’s intrinsic properties are tunable by changing the activity level.

### 5.1 Muscle response to perturbations at dynamic-scenario

During dynamic-scenario experiments, muscle fibers initially react elastically to the sudden perturbation, known as short-range stiffness (Kirsch et al. 1994; Rack and Westbury 1974). Muscles then transition into a viscoelastic behavior (Fig. 3A). We found no significant changes in fiber work loops, although different perturbations changed stretch velocities (Fig. 6C). We calculated fiber work starting at the beginning of the stretch, until the end of the short-range stiffness phase at 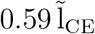 (Fig. 3A). Our observation is in agreement with the reported constant short-range stiffness for stretch velocity ranges similar to ours (3 L_0_/s to 5 L_0_/s) (Pinniger et al. 2006; Rack and Westbury 1974).

After the short-range stiffness phase, i.e., from 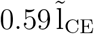 to the end of preflex, the force response became nonlinear, and velocity adaptation occurred. In this phase, higher stretch velocities cause higher forces (Fig. 3A) and preflex work (Fig. 4A). Both observations are in agreement with the reported work increase with increasing stretch velocity for the eccentric phase of ramp-like stretch-shortening cycles (Tomalka et al. 2021).

Two factors contribute to the increasing preflex work in our study. The first factor is the higher force. However, suppose the same stretch is considered for calculating the preflex work (Fig. 5A, inset). In that case, differences in preflex work between perturbations are insignificant. Second, higher velocities cause a larger fibers stretch in the preflex phase (Fig. 3A). If the larger stretch is fully considered, differences in preflex work between the perturbation cases become significant (Fig. 5). For the specified area, all energetic differences between perturbations are the result of muscle fibers’ viscous behavior since stretch amounts are identical. We find a rising trend in dissipated energy with the increase of stretch velocity from step up to step down perturbations. Hence, muscles adapt their work response primarily due to a change of stretch caused by locomotion perturbation.

### 5.2 Tuning the force and energy response by activity level

The increased muscle activity strategy increases walking robustness (Haeufle et al. 2018). To test whether increased activity leads to higher muscle stiffness and work in this scenario, we conducted muscle fiber experiments with activity levels of 5%, 15% and 25% for each perturbation case while keeping the kinematics identical.

The results show that the short-range stiffness increases with activity level (Fig. 6A), which is in line with previous findings (Campbell et al. 2014). As short-range stiffness is likely due to the stretch of attached cross-bridges (Getz et al. 1998; Pinniger et al. 2006), an increase in short-range stiffness can be explained by the increasing number of attached cross-bridges with increasing activity level (Metzger and Moss 1990).

Additionally, we found that higher activity levels result in significantly higher preflex work (Fig. 4A). Tomalka et al. (2021) likewise reported an increase in work with an increasing number of active cross-bridges in the eccentric phase of the ramp-like stretch-shortening cycles. An increase in preflex work with increasing activity level might be explained by an increasing number of forcibly detached cross-bridges after the short-range stiffness phase. The number of forcibly detached cross-bridges might increase (at the given stretch kinematics) as a fraction of the increasing number of attached cross-bridges with activity level (Wahr and Rall 1997). Forced detachment of cross-bridges is expected (Weidner et al. 2022) in the range of the tested stretch velocities (3 l_opt_/s to 5 l_opt_/s; Fig. 6). Additionally, viscoelastic properties of non-cross-bridge structures (e.g., titin) might contribute to energy dissipation in a velocity-dependent manner (Freundt and Linke 2019; Herzog et al. 2014; Tomalka et al. 2021). Furthermore, with higher activity, the differences between perturbation cases become more prominent, and, for 25%, even significant (Fig. 4A).

Consequently, humans and animals may tune their muscle stiffness during the short-range stiffness phase (Fig. 6A) and work afterward (Fig. 5A) utilizing the activity level. Thus, increased muscle stiffness and work in preparation for an expected perturbation are possible by increasing the muscle pre-activity level.

### 5.3 Dynamic-versus quasistatic-scenario

During the preflex phase of the quasistatic-scenario experiments, when muscles are stretched with negligible velocities, they respond with a linear force increase which is almost the same regardless of the perturbation case (Fig. 3B). Comparison of dynamic-scenario and quasistatic-scenario experiments show that velocity is not only adding a viscous behavior to the response (Fig. 3A vs. B), but also a visible short-range stiffness contribution (Fig. 6). In previous iso-velocity stretch experiments, the initial force response (i.e., short-range stiffness) was velocity dependent (Pinniger et al. 2006; Rack and Westbury 1974), especially when the strain rate was varied over several orders of magnitude. For example, Pinniger et al. (2006) showed that the initial force response differs between slow (0.1 L_0_/s) and fast (2 L_0_/s) stretches. However, for contractions faster than 2 L_0_/s they observed no significant difference in the short-range stiffness. Weidner et al. (2022) observed differences in the short-range stiffness between 0.01 L_0_/s and 1 L_0_/s stretches. Hence, even though the short-range stiffness is independent of velocity at the range we worked in dynamic-scenario experiments (3 L_0_/s to 5 L_0_/s, Fig. 6A), the difference between actual and quasistatic-scenario experiments performed at a maximum of 0.05 L_0_/s showed that the short-range stiffness represents a velocity-dependent muscle behavior.

Interestingly, even though the dynamic behaviors during the preflex phase differ between quasistatic-scenario and dynamic-scenario conditions (Fig. 3A,B), we observed almost equal amounts of mechanical work at quasistatic-scenario compared to dynamic-scenario stretches for each activity level (Fig. 4A,B).

Possibly, myosin heads are detached forcibly from actin at high velocities during eccentric contractions. This will decrease muscle force generation (Weidner et al. 2022) observed during in-situ and in-vitro experiments (Fukutani et al. 2019; Griffiths et al. 1980; Krylow and Sandercock 1997; Till et al. 2008; Tomalka et al. 2020; Weidner et al. 2022). On the other hand, during the quasistatic-scenario stretches, ultra-slow-speed stretches allow cross-bridges to bind easier and longer than during rapid contractions (Herzog 2018; Huxley 1957). Hence, similar forces during the preflex stretch phase result in similar amounts of energy dissipation in quasistatic-scenario and dynamic-scenario experiments.

### 5.4 Muscle fibers versus Hill-type muscle model

Since Hill’s empirical investigations of muscle contraction dynamics, Hill-type models have played a crucial role in biomechanics research (Hill 1938; Rode and Siebert 2017). These models have been improved over the years, but they are still limited in predicting muscle forces, especially during eccentric (lengthening) contractions (Siebert et al. 2021; Till et al. 2008). Surprisingly, our results show that the magnitude and trends in mechanical work predicted by the Hill-type contractile element model are similar to the work of muscle fibers for fast eccentric contractions (Fig. 4A,C). This is an unexpected outcome since the force response of the Hill-type muscle model and muscle fibers differs (Fig. 3A vs. C). We show that the main source of preflex-work adaptation to perturbation height is the amount of muscle stretch rather than the viscous force adaptation.

Our quasistatic-scenario experiments and simulations proved that the force-length relation of the Hill-type muscle model could accurately estimate the length-dependent force and mechanical work response of muscle fibers for the investigated conditions (relatively short muscle fibers at low activity levels). Because the length-dependent behavior of muscle fibers is the main force contributor during preflex, the Hill-type muscle model predicts work for the larger stretch in response to the fast contraction reasonably well.

Although the Hill-type muscle model estimates work during preflex with good accuracy, it still requires improvements for better force prediction during fast contractions (Fig. 3A,C). The shortrange stiffness had previously been observed in other fiber experiments (Rack and Westbury 1974; Tomalka et al. 2021; Weidner et al. 2022). We observed that the short-range stiffness was activity- and velocity-dependent (Fig. 6), at least for the velocity difference between dynamic-scenario and quasistatic-scenario experiments (Fig. 3A,B). However, our Hill-type muscle model cannot generate the high-stiffness response of a short-range stiffness, since short-range stiffness is not a built-in mechanical property (Haeufle et al. 2014). So far, only a few studies included modeling short-range stiffness (Cui et al. 2008). De Groote et al. (2017) applied a muscle model with a short-range stiffness in a locomotion study and showed its functional relevance in terms of a reduction of unrealistic joint angles and improved locomotion stability during simulated gaits, compared to classic Hill-type muscle models. Together with our results, we expect that Hill-type muscle models that feature short-range stiffness should provide a better force estimation at and immediately after impact. Therefore, we consider short-range stiffness an essential model feature for the understanding of gait mechanics leading to stable locomotion.

The force adaptation to the perturbation velocity after the short-range stiffness (Fig. 3A vs. C) is also not accurately modeled in the Hill-type muscle model. Here, the Hill-type muscle model operates in the plateau region of the eccentric force-velocity relation (Fig. 2). The model, therefore, does not show any adaptation of the force due to the perturbation-related changes in fiber velocity, in contrast to the observations in the experiments (Fig. 3A vs. C). This plateau-form of the eccentric force-length relation was introduced by Soest and Bobbert (1993), while their results are consistent with our simulation data, they do not explain the experimental results of the present study. Possible reasons for the deviation of the experiment can be the starting length of the contraction, the sub-maximal activity level or the underlying model. However, the results of Krylow and Sandercock (1997) suggest that the starting length has no influence on the point of occurrence of the eccentric force-velocity-relation’s plateau. Regarding the sub-maximal activity level and its effects, it is known that the calcium concentration has an influence on the cross-bridge kinetics (Brenner 1988). Nevertheless, to the best of our knowledge, there is no study there is no study that looked at the dependency of the force-velocity relation on the activity level. A likely explanation for the discrepancy is simplification of the contraction by the model. For example, the Hill-type muscle model lacks the implementation of “Give” (Flitney and Hirst 1978), which on the one hand is speed-dependent and on the other hand occurs directly after the end of the short-range stiffness (Weidner et al. 2022). The force and work responses show that the modeling of eccentric muscle behavior needs to be modified for better estimation of the dynamic response to perturbations during fast eccentric contractions.

### 5.5 Study limitations

This study aimed at analyzing how a single muscle fiber reacts to ground perturbations in real life. We conducted single-leg hopping simulations using a Hill-type muscle model as a knee extensor muscle to generate kinematic boundary conditions for in-vitro experiments. However, Hill-type muscle models have limitations discussed in previous chapters, and simulation and real-life muscle lengthening may differ. Additionally, our in-vitro experimental setup allows only constant activity levels. Even though in the hopping simulations after the preflex phase activity rises, due to the setup limitations, we perform the kinematic analysis with the constant preflex activity level, which is not the case in in-vivo locomotion (Müller et al. 2015). Thus, we only focused our analysis on the preflex phase. Since we do not consider a full work loop, this study’s design does not directly allow us to calculate damping, i.e., the amount of energy dissipation in a full cycle. However, the velocity-dependent adaptation of preflex work indicates a viscous-like response, which we identify as a damping behavior.

## 6 Conclusion

Previous experimental and simulation studies indicated that muscles’ preflex capability to adapt force to unexpected ground conditions is essential in stabilizing locomotion. Our study confirms these findings and shows three mechanisms: (1) muscle force adapts to the change in stretch velocity caused by a perturbation; (2) the overall fiber stretch in the preflex duration is larger for larger stretch velocities resulting in increased preflex work; (3) with increasing muscle activity short-range stiffness and muscle force increase. Mechanism (1) is the hypothesized viscous effect of the force-velocity relationship but plays a minor role compared to the mechanism (2). Together, (1) and (2) result in a beneficial and significant adaptation of muscle force to perturbations and thus confirm the preflex hypothesis. Mechanism (3) allows for a simple neuronal strategy to tune the muscle properties to ground conditions and unexpected perturbations and aligns with feed-forward strategies observed in human locomotion (Müller et al. 2012).

## Author Contributions

M.A., S.W., A.B.-S., T.S. and D.H. conceptualized the project. F.I. and M.A. conducted computer simulations. M.A. and S.W. conducted the experiments and analyzed the data. M.A. and S.W. prepared the manuscript, and all authors revised and approved the submitted version.

