## Supplementary material for "Muscle Preflex Response to Perturbations in locomotion: In-vitro experiments and simulations with realistic boundary conditions": Fig. S1

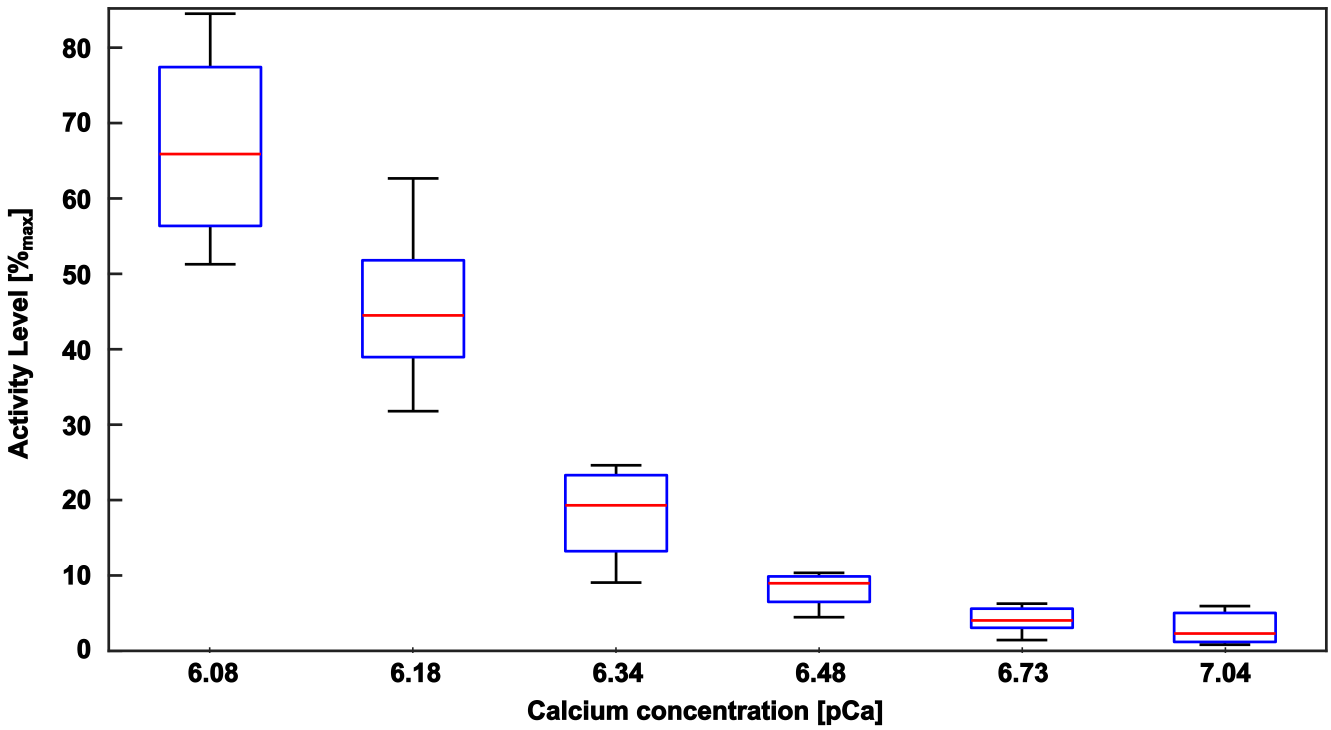


Figure S1: Measured activity level of skinned fibers (n=7) from the muscle used in the current study depending on the pCa of the experimental solution. Temperature while testing was 12 °C as in the perturbation experiment.
