## Supplementary material for "Muscle Preflex Response to Perturbations in locomotion: In-vitro experiments and simulations with realistic boundary conditions": Fig. S2

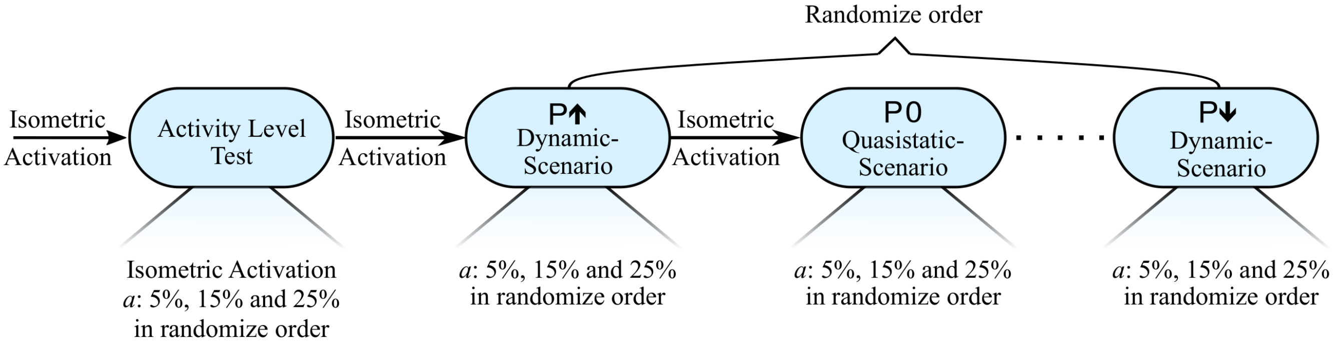


Figure S2: The flow chart of an experimental day is shown here. First the activity level for every fiber was checked using the 6.73 , 6.34 and 6.3 pCa concentration solution to ensure that the experiment is matching the simulation condition. Afterwards the experimental blocks were conducted. One block contained all contractions (n=3) of a perturbation for one velocity-scenario. The order of the blocks were randomized on the day of the experiment. Between two blocks a reference contraction at optimal length and full activity was conducted to check for the degradation of the skinned fiber.
