## Supplementary material for "Muscle Preflex Response to Perturbations in locomotion: In-vitro experiments and simulations with realistic boundary conditions": Fig. S3

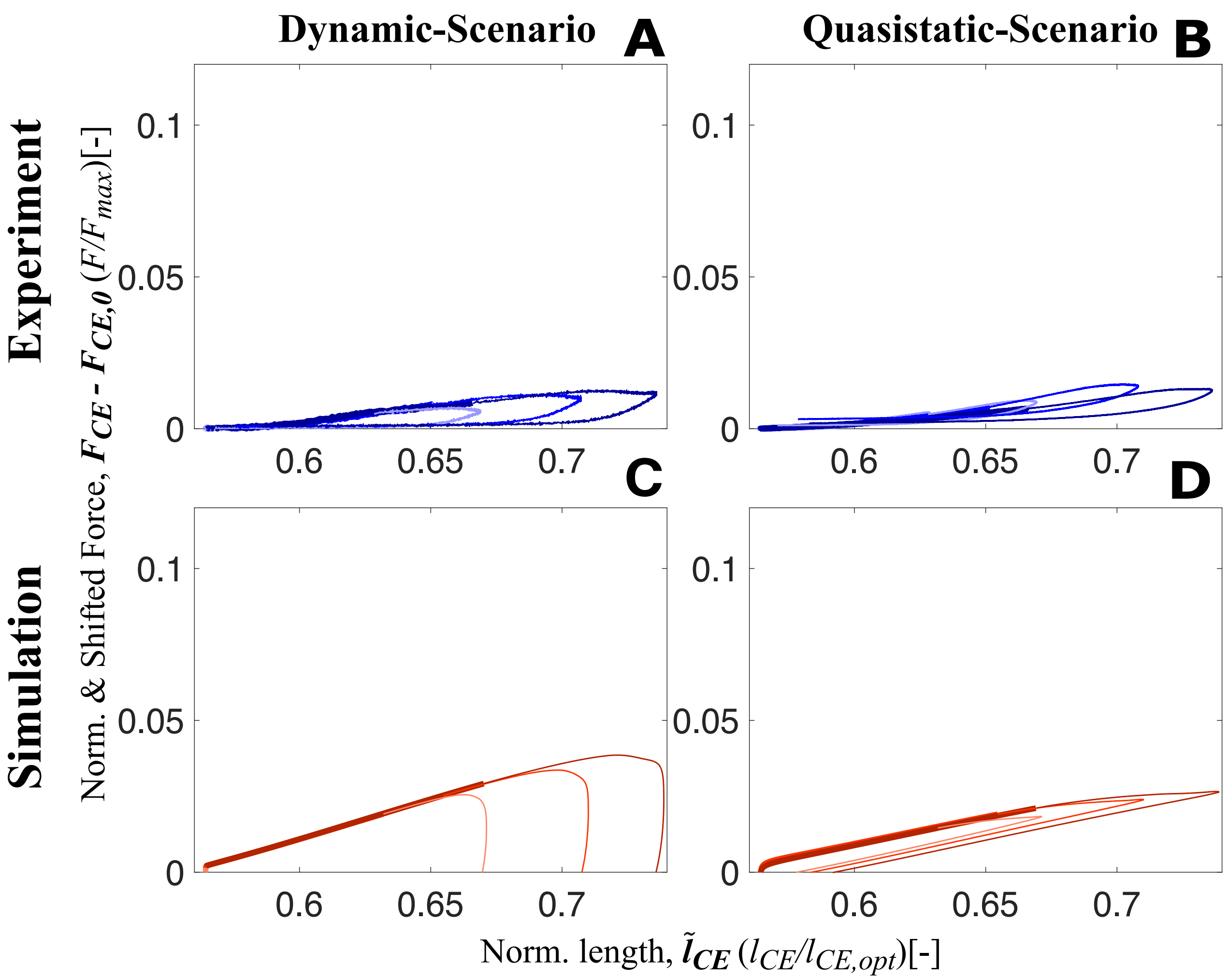


Figure S3: Shifted work loops for dynamic-scenario and quasistatic-scenario analysis step up(P↑), no (P0) and step down (P↓) perturbations for both experiments (**A-B**) and simulations (**C-D**) at 5 % activity level. The experimental data presented on **A** and **B** show the mean of all experimental trials. From touch-down to toe-off, all stretch-shortening cycle loops are plotted in the clockwise direction, and the thick and thin sections of the loops represent the preflex and remaining part of the stretch-shortening cycle, respectively.
