## Supplementary material for "Muscle Preflex Response to Perturbations in locomotion: In-vitro experiments and simulations with realistic boundary conditions": Fig. S6

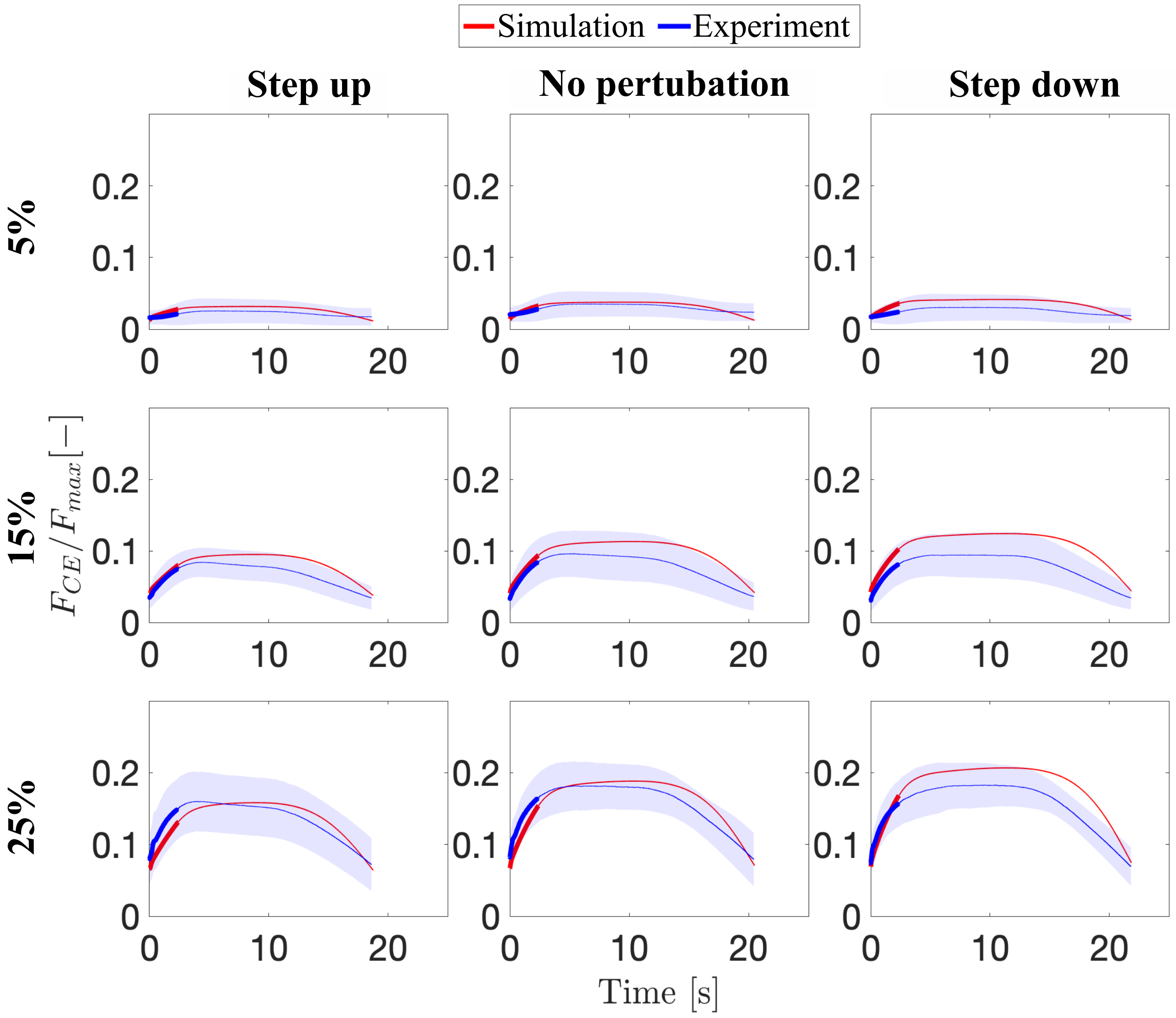


Figure S6: Force generated in quasistatic-scenario by both muscle fibers and the Hill-type muscle model during one hopping cycle — from touch-down to toe-off — are presented for all perturbation and activity levels. The thick and thin sections of the loops represent the preflex and remaining part of the stretch-shortening cycle, respectively. the blue line and shaded area represent the mean of all experimental trials and the standard deviation, respectively.
