## Supplementary material for "Muscle Preflex Response to Perturbations in locomotion: In-vitro experiments and simulations with realistic boundary conditions": Tab. S1

Table S1: Statistical comparison of muscle fibers' perturbation response for each activity level. Significantly different results are indicated by *

| Condition | $a$ | Parameter | $\chi^2$ | $p$ | P0 vs. P↑($r$) | P0 vs. P↓($r$) | P↓ vs. P↑ ($r$) |
|---|---|---|---|---|---|---|---|
| Dynamic-Scenario | 5% | Preflex work | 8.222 | 0.016* | 0.029(.41)* | 1 | 0.055 |
| | | SRS | 0.889 | 0.641 | - | - | - |
| | | Work Fig. 5A | 1.556 | 0.459 | - | - | - |
| | 15% | Preflex work | 16.222 | 0.001* | 0.055 | 0.297 | 0.001(0.63)* |
| | | SRS | 0 | 1 | - | - | - |
| | | Work Fig. 5A | 4.667 | 0.097 | - | - | - |
| | 25% | Preflex work | 14.889 | 0.001* | 0.029(0.41)* | 0.716 | 0.001(0.59)* |
| | | SRS | 1.556 | 0.459 | - | - | - |
| | | Work Fig. 5A | 3.556 | 0.169 | - | - | - |
| Quasistatic-Scenario | 5% | Preflex work | 4.222 | 0.121 | - | - | - |
| | | Stiffness | 4.222 | 0.121 | - | - | - |
| | 15% | Preflex work | 16.222 | 0.001* | 0.055 | 0.297 | 0.001(0.63)* |
| | | Stiffness | 0.667 | 0.717 | - | - | - |
| | 25% | Preflex work | 16.222 | 0.001* | 0.055 | 0.297 | 0.001(0.63)* |
| | | Stiffness | 0.667 | 0.717 | - | - | - |
| Dynamic vs. Quasistatic-Scenario Fig. 5B | 5% | Preflex work | 2.667 | 0.264 | - | - | - |
| | 15% | Preflex work | 4.222 | 0.121 | - | - | - |
| | 25% | Preflex Work | 1.556 | 0.459 | - | - | - |
