## Supplementary material for "Muscle Preflex Response to Perturbations in locomotion: In-vitro experiments and simulations with realistic boundary conditions": Tab. S2

Table S2: Statistical comparison of activity differences for each perturbation case. Significantly different results are indicated by *

| Condition | P | Parameter | $\chi^2$ | p | 05% vs. 15% | 05% vs. 25% | 15% vs. 25% |
|---|---|---|---|---|---|---|---|
| Dynamic-Scenario | P0 | Preflex work | 18 | 0.001* | 0.102 | 0.001(0, 67)* | 0.102 |
| | | SRS | 18 | 0.001* | 0.102 | 0.001(0.67)* | 0.102 |
| | | Work Fig. 5A | 12.667 | 0.002* | 0.472 | 0.001(0.56)* | 0.102 |
| | P↑ | Preflex work | 18 | 0.001* | 0.102 | 0.001(0.67)* | 0.102 |
| | | SRS | 18 | 0.001* | 0.102 | 0.001(0.67)* | 0.102 |
| | | Work Fig. 5A | 18 | 0.001* | 0.102 | 0.001(0.67)* | 0.102 |
| | P↓ | Preflex work | 18 | 0.001* | 0.102 | 0.001(0.67)* | 0.102 |
| | | SRS | 18 | 0.001* | 0.102 | 0.001(0.67)* | 0.102 |
| | | Work Fig. 5A | 18 | 0.001* | 0.102 | 0.001(0.67)* | 0.102 |
| Quasistatic-Scenario | P0 | Preflex work | 16.22 | 0.001* | 0.055 | 0.001(0.63)* | 0.297 |
| | | Stiffness | 12.667 | 0.002* | 0.102 | 0.001(0.56)* | 0.472 |
| | P↑ | Preflex work | 16.22 | 0.001* | 0.055 | 0.001(0.63)* | 0.297 |
| | | Stiffness | 11.556 | 0.003* | 0.055 | 0.003(0.52)* | 1 |
| | P↓ | Preflex work | 18 | 0.001* | 0.102 | 0.001(0.67)* | 0.102 |
| | | Stiffness | 11.556 | 0.003* | 0.055 | 0.003(0.52)* | 1 |
| Dynamics vs. Quasistatic-Scenario Fig. 5B | P0 | Preflex work | 2 | 0.368 | - | - | - |
| | P↑ | Preflex work | 0.222 | 0.895 | - | - | - |
| | P↓ | Preflex work | 0.222 | 0.895 | - | - | - |
