## Supplementary material for "Muscle Preflex Response to Perturbations in locomotion: In-vitro experiments and simulations with realistic boundary conditions": Tab. S3

Table S3: Statistical comparison between Dynamic and Quasistatic Scenario. Significantly different results are indicated by *

| Activity Level | Parameter | P | $z$ | $p$ | $r$ |
|---|---|---|---|---|---|
| 5% | Preflex work | P0 | 0.533 | 0.594 | - |
| | | P↑ | 0.296 | 0.767 | - |
| | | P↓ | 1.333 | 0.182 | - |
| | SRS | P0 | 0 | 1 | - |
| | | P↑ | 0 | 1 | - |
| | | P↓ | 0 | 1 | - |
| 15% | Preflex work | P0 | 0.667 | 0.505 | - |
| | | P↑ | 0.533 | 0.594 | - |
| | | P↓ | 0.667 | 0.505 | - |
| | SRS | P0 | 1.333 | 0.182 | - |
| | | P↑ | 2 | 0.046* | 0.67 |
| | | P↓ | 2.666 | 0.008* | 0.89 |
| 25% | Preflex work | P0 | 0.667 | 0.505 | - |
| | | P↑ | 0.652 | 0.515 | |
| | | P↓ | 0.667 | 0.505 | - |
| | SRS | P0 | 2.666 | 0.008* | 0.89 |
| | | P↑ | 2.666 | 0.008* | 0.89 |
| | | P↓ | 2 | 0.046* | 0.67 |
